# Semaphorin 4D induced inhibitory synaptogenesis restores benzodiazepine sensitivity in a mouse model of Status Epilepticus

**DOI:** 10.1101/2022.10.25.513689

**Authors:** Susannah S. Adel, Aidan Evans-Strong, Vernon R.J. Clarke, Jamie Maguire, Suzanne Paradis

**Author notes:** co-corresponding authors to whom correspondence should be addressed **<u>AUTHOR INFORMATION:</u> Co-corresponding authors:** Suzanne Paradis, Department of Biology, Brandeis University, 415 South Street, Waltham, MA 02454, Jamie Maguire, Neuroscience Department, Tufts University School of Medicine, 136 Harrison Avenue, Boston, MA 02111. **All other authors:**. Susannah S. Adel, Department of Biology, Brandeis University, 415 South Street, Waltham, MA 02454. Vernon Clarke, Department of Biology, Brandeis University, 415 South Street, Waltham, MA 02454. Aidan Evans-Strong, Neuroscience Department, Tufts University School of Medicine, 136 Harrison Avenue, Boston, MA 02111.

## Abstract

Previously we demonstrated that intra-hippocampal infusion of purified, Semaphorin 4D (Sema4D) extracellular domain into the mouse hippocampus rapidly promotes formation of GABAergic synapses and decreases seizure susceptibility in mice. Given the relatively fast action of Sema4D treatment revealed by these studies, we sought to determine the time course of Sema4D treatment on hippocampal network activity using an acute hippocampal slice preparation. We performed long-term extracellular recordings from area CA1 encompassing a 2 hour application of Sema4D and found that hippocampal excitation is suppressed hours following treatment. We also asked if Sema4D treatment could ameliorate seizures in an acute seizure model: the kainic acid (KA) mouse model of Status Epilepticus (SE). We demonstrate that Sema4D treatment suppresses seizures and lessens benzodiazepine (BZD) insensitivity which is a hallmark of refractory SE. Lastly, we sought to explore alternative methods of Sema4D delivery to hippocampus and thus created an Adeno Associated Virus expressing the extracellular domain of Sema4D. Our data reveal that virally delivered, chronically overexpressed Sema4D extracellular domain promotes GABAergic synapse formation and suppresses seizure activity similar to the actions of purified Sema4D protein. These results provide proof of concept that Sema4D-mediated GABAergic synapse development could combat BZD insensitivity in intractable forms of epilepsy, and that viral delivery of Sema4D is an efficacious and promising delivery method.

## 1. Introduction

Status epilepticus (SE) is a life-threatening neurological emergency characterized by continuous seizure activity lasting greater than 5 minutes (Trinka et al., 2015) which can have serious long-term consequences including neuronal injury and death. First line treatment for SE is intravenous or intramuscular administration of benzodiazepines (BZD; e.g. diazepam) (Cruickshank et al., 2022). Benzodiazepines enhance the activity of GABA_A_ receptor subunits, thereby affecting existing GABAergic synapses, and increasing inhibitory tone in the brain. Unfortunately, ∼30% of SE patients do not respond to treatment with BZD plus at least one other anti-seizure medication, resulting in refractory SE (RSE) which has a mortality rate of approximately 35% (Mayer et al., 2002; Shorvon & Ferlisi, 2012). Thus, the need for innovative anti-seizure medications to treat SE and RSE is dire.

While the cellular pathophysiology underlying BZD resistance in RSE remains unclear, current models to explain this event include: aberrant removal of GABA_A_Rs from the neuronal cell surface (Naylor, 2005), increased AMPA and NMDA receptor abundance at synapses (Loscher, 2015; Wasterlain & Chen, 2008), and dysfunction of the type-2 K^+^-Cl^-^ cotransporter (KCC2) causing elevated intracellular Cl-concentrations, leading to depolarizing action of GABA_A_Rs (reviewed in (Deeb, Maguire, & Moss, 2012). We and others demonstrated that the extracellular domain of Sema4D promotes the stabilization and development of GABAergic synapses on a rapid time scale (∼30 minutes) (Acker, Wong, Kang, & Paradis, 2018; Frias et al., 2019; Kuzirian, Moore, Staudenmaier, Friedel, & Paradis, 2013); these synapses become functional within two hours (Kuzirian et al., 2013). We reasoned that if maladaptive GABA_A_R internalization makes a significant contribution to the development of RSE, treatment with the pro-synaptogenic molecule Sema4D could impart therapeutic effects in mouse models of this disease.

In addition, our previous studies using *in vivo* seizure models (intravenous pentylenetetrazol and electrical kindling) demonstrated that acute infusion of the purified Sema4D extracellular domain (Sema4D-Fc) into hippocampus increases GABAergic synapse density, increases time to seizure onset, and decreases seizure severity (Acker et al., 2018). Based on these data we proposed a model whereby Sema4D treatment in hippocampus promotes GABAergic synapse formation, thus increasing inhibitory tone and suppressing seizures (Acker et al., 2018).

However, whether and how Sema4D treatment directly affects hippocampal circuit activity, in addition to the time course of this effect, remain unanswered questions. Therefore, we used extracellular recordings to assay population spike amplitude in the CA1 region of acute hippocampal slice before, during, and after 2hrs of Sema4D treatment (∼6 hrs total for entire recording). In addition, the proposed mechanisms of Sema4D-dependent seizure suppression led us to hypothesize that Sema4D delivery to the intact hippocampus could be mechanistically complementary to that of BZD and increase BZD efficacy in combating seizure. We also sought to determine if chronic delivery of Sema4D protein via viral mediated gene transduction in the hippocampus, a more generalizable and therapeutically tractable delivery method, could suppress seizure onset and severity.

We elected to test these questions in the KA model of SE to determine Sema4D’s ability to counteract BZD insensitivity when Sema4D is administered in conjunction with BZD. KA-induced SE is an established rodent model of temporal lobe epilepsy which progresses to SE and development of pharmacoresistance (reviewed in Reddy & Kuruba, 2013). We delivered purified Sema4D-Fc protein via cannula to the hippocampus of mice and subjected them to KA-induced SE and subsequent BZD treatment while monitoring seizure severity using EEG. Next, we developed and validated adeno associated virus expressing Sema4D-ECD and tested whether chronic Sema4D-ECD expression via viral mediated gene transduction could increase GABAergic synapse density. Finally, we asked if viral delivery of Sema4D to intact hippocampus could suppress seizures and/or increase efficacy of BZD treatment in the KA model of SE.

## 2. Methods

### 2.1 Animals

C57BL/6 male mice and adult Long-Evans male and female rats were purchased from the Charles River Laboratories and housed in the animal facility at Brandeis University or at Tufts University. Animals were maintained with a 12-hour light-dark cycle. Food and water were available ad libitum. Animal procedures were performed with approval from the Brandeis University Institutional Animal Care and Use Committee and the Tufts University School of Medicine Animal Care and Use Committee in accordance with the Guide for the Care and Use of Laboratory Animals (NRC). Animal studies were performed in compliance with ARRIVE guidelines as described in this methods section.

### 2.2 Population spike recordings

Brains were isolated from Long-Evans rats of both sexes between 20-25 weeks old. The brain was rapidly dissected and placed in ice-chilled, oxygenated modified artificial cerebrospinal fluid (aCSF) comprising of (in mM): NaCl 124, KCl 3.7, NaHCO_3_ 24.6, CaCl_2_ 1, MgSO_4_ 3, *D*-glucose 10, KH_2_PO_4_ 1.2 saturated with 95% O_2_ / 5% CO_2_. Parasagittal slices (400 μm thick) containing the hippocampal region were prepared in ice-chilled, oxygenated aCSF using a Leica VT1200 vibrating microtome (Leica Biosystems Inc.).

Following equilibration for at least an hour, slices were transferred to a brain slice interface chamber (model BSC2, Scientific Systems Design, Inc.); slices rested on filter paper at the interface of the perfusing solution (0.4 ml min^-1^) which comprised of standard aCSF saturated with 95% O_2_ / 5% CO_2_. Field recordings were obtained using a glass microelectrode (resistance ∼ 2-4MΩ) containing 3M NaCl placed at the stratum pyramidale/stratum oriens border. Responses were evoked by stimulating two electrodes placed within the Schaffer-collateral-commissural fibers. Baseline stimulation was every 150 seconds for each input with a 75 second interval between alternating inputs. Independence of inputs was assessed using a paired-pulse protocol. Stimulation intensity was adjusted such that the baseline population spike amplitude was approximately 40% of the maximum amplitude. Recordings were filtered at 3–10 kHz using an Axoclamp 2A Amplifier (Molecular Devices), and collected for online analysis at a sampling rate of 20kHz (Nguyen, 2013; Redondo et al., 2010) using WinLTP software (www.winltp.com; (Anderson & Collingridge, 2007).

### 2.3 Electroencephalogram (EEG) recording

EEG recordings from male C57BL/6 mice were carried out as previously described (Hooper, Fuller, & Maguire, 2018; Sivakumaran & Maguire, 2016). Mice (10-12 weeks of age) were anesthetized with 100 mg/kg ketamine and 10 mg/kg xylazine and a prefabricated headmount (part # 8201; Pinnacle Technology, Inc) was affixed to the skull with four screws and dental cement. The screws serve as differential EEG leads, two of which were placed bilaterally anterior and two posterior to bregma. The animals were allowed to recover for a minimum of 5 days before experimentation EEG recordings and analyses were performed by two different experimenters with blinding to condition.

EEG recordings were collected in awake, behaving animals using a 100x gain preamplifier high pass filtered at 1.0 Hz (Pinnacle Technology, part #8202-SE) and tethered turnkey system (Pinnacle Technology, part #8200). Seizure susceptibility was assessed in response to the chemoconvulsant, KA (20mg/kg, i.p.) and electrographic seizure activity was recorded for 1hr before and 2hrs following KA administration and before diazepam administration. In brief, epileptiform activity was considered to be paroxysmal activity having a sudden onset and an amplitude at least 2.5x the standard deviation of the baseline and a consistent change in the Power of the fast Fourier transform of the EEG. In the KA model, the animals typically enter SE by 1 hour post-KA administration; SE was defined as persistent, unremitting epileptiform activity lasting longer than 5 consecutive minutes. Our definition of “epileptiform activity” includes both discrete ictal events and periods of SE. These criteria have been used previously by our group (O’Toole, Hooper, Wakefield, & Maguire, 2014,) (Maguire, Stell, Rafizadeh, & Mody, 2005) (Lee & Maguire, 2013) as well as by experts in the field (Castro et al., 2012). Seizure latency was defined as the time elapsed from the KA injection to the start of the first electrographic seizure. The % time epileptiform activity was calculated as the cumulative time of all epileptiform activity during a 60-min recording period divided by 60 min. The latency to SE was calculated as the time elapsed from the KA injection to the start of the SE (first 5 mins of unremitting epileptiform activity).

For the Sema4D infusion experiments, mice were implanted with a guide cannula into the dorsal hippocampus (A/P: -2.0mm; M/L: ± 1.5mm; D/V: -2mm) during EEG headmount attachment and Sema4D (100nM, 500nl) or vehicle (0.9% sterile injection saline, 500nl) was infused into the hippocampus 1 hr prior to KA administration. For virally delivered Sema4D, 500nl of AAV-Sema4D (2.55 Gc/mL) or control virus was stereotaxically injected into the dorsal hippocampus 1 week prior to KA administration. To assess the ability of Sema4D to restore BZD sensitivity, we administered 5mg/kg diazepam 2hrs after KA administration as previously described by our laboratory (Sivakumaran & Maguire, 2016). The diazepam sensitivity (as measured by a reduction in the power of the electrographic signal and cessation of epileptiform activity) was recorded as previously described (Hooper et al., 2018; Sivakumaran & Maguire, 2016). Mice were considered to be pharmacoresistant if diazepam treatment failed to suppress epileptiform activity within 10 mins of administration.

### 2.4 Immunohistochemistry and analysis for virus-injected animals

Male C57BL/6 mice were injected with AAV-Sema4D-ECD or AAV-CD4-ECD virus as above and sacrificed 2-3 weeks post-injection. The brains were removed and fixed in 4% paraformaldehyde for 24 hours, transferred to 10% sucrose for 24 hours, and then transferred to 30% sucrose for 24 hours. They were then embedded in Optimal Cutting Temperature (OCT) compound and frozen down for storage at - 80°C. The brains were sectioned at 40 μm using a cryostat (Leica) and slices were suspended in PBS at 4°C until being processed for immunohistochemistry.

Representative slices from the hippocampus were chosen for staining. The sections were blocked in PBS with 10% normal goat serum (NGS) and 0.3% Triton for one hour at room temperature prior to incubation with the primary antibody mouse anti-GAD65 (EMD Millipore, MAB3551) at a concentration of 1:100 overnight at 4°C. The sections were then incubated in the secondary antibody (anti-Ms Alexa 488, Invitrogen, A32766) at 1:200 for two hours at room temperature and mounted with Vectashield Hardset antifade mounting medium with DAPI (Vector Laboratories, H-1500). Imaging was performed on a Leica SP8 Confocal microscope using a 40x oil objective. 2-3 images per hemisphere per section were taken of the hippocampus near A/P - 2.0 (the virus injection site). Microscope and laser settings were kept constant across all images. Quantification of GAD65 immunofluorescence was performed using ImageJ. An outline was traced around the CA1 region of the hippocampus and the fluorescence intensity in the region of interest was quantified. For each image, the CA1 mean intensity was normalized to the mean intensity of background.

### 2.5 Organotypic slice culture

C57BL/6 mouse brains were dissected from P6-P8 animals of both sexes into cutting solution (126mM NaCl, 25mM NaHCO_3_, 3mM KCl, 1mM NaH_2_PO_4_, 20mM dextrose, 2mM CaCl_2_, 2mM MgCl_2_ in deionized water at 315-319 mOsm) (Stoppini, Buchs, & Muller, 1991). Coronal slices were taken at a thickness of 300 μm using a tissue chopper (Compresstome VF-200, Precisionary Instruments Inc.). Individual slices were placed on cell culture inserts (0.4 um pore size, Millipore). Organotypic culture media (2mM glutamax, 1mM CaCl_2_, 2mM MgSO_4_, 12.9mM d-glucose, 0.08% ascorbic acid, 18 mM NaHCO_3_, 35mM HEPES, 20% horse serum, 1 mg/mL insulin in minimum essential media) at pH 7.45 and 305 mOsm was added outside of the inserts. Slices were maintained for 6 days in vitro at 35°C and 5% CO_2_ with media replacements every other day.

One day after harvesting slices, a solution containing AAV9.hsyn viruses encoding either full-length or the extracellular domain of either Sema4D or CD4 (designed and purified by Vector Biolabs) in combination with the same amount of AAV9.hsyn.GFP virus (Addgene, #105539-AAV9) was pipetted onto the hippocampus within the slice (1 μL on each hemisphere, each virus in the solution at 2.55 × 10^12^ Gc/mL). A subset of slices were infected with only AAV9.hsyn.GFP and at DIV6 were treated with human Sema4D-Fc ectodomain/human Fc fusion protein (Sema4D-Fc; R&D Systems, #7470-S4) or Fc control protein (R&D Systems, #110-HG) at a final concentration of 2 nM/well for 2 hours before being fixed immediately following.

### 2.6 Immunofluorescence performed in organotypic slice cultures

To quantify synapse density at DIV6, organotypic slices were fixed in 4% paraformaldehyde/4% sucrose for 20 minutes at 4°C. After 3 × 10-minute washes in PBS, slices were incubated overnight in permeabilization solution (0.1% Triton-X in PBS) followed by an overnight incubation in blocking solution (20% bovine serum albumin with 0.1% Triton-X in PBS) at 4°C. Next, slices were incubated in primary antibody anti-GAD65 (EMD Millipore, #MAB351) at 1:150 in blocking solution and incubated overnight at 4°C followed by 3 × 10-minute PBS washes and incubation with secondary antibody anti-mouse-Cy3 (Jackson Laboratories, #115-165-003) at 1:500 into secondary solution (blocking solution diluted 1:1000 in PBS) for 2 hours at room temperature. Following 3 × 10-minute PBS washes, slices were mounted (insert side down) on slides in Vectashield + DAPI mounting media (Vector Laboratories).

### 2.7 Imaging and analysis of organotypic slice cultures

16-bit images of neurons were acquired on a Zeiss LSM880 Confocal microscope using a Plan-Apochromat 63x/1.40 Oil DIC M27 objective. Within each experiment (slices collected on the same day), images were acquired with identical settings for laser power, detector gain, and amplifier offset. Settings were initially optimized across multiple control slices to avoid oversaturation. Images were acquired as z-stacks (5-15 optical sections, 0.5 μm step size) for each of 4-5 fields of view per hemisphere (134.95 μm x 134.95 μm) containing the pyramidal cell layer in CA1 from each slice. Within each z-stack, the image with the highest fluorescent intensity for GAD65 signal was chosen for analysis and neurons were selected at random for analysis using only the GFP signal, thus blinding the experimenter to the GAD65 signal in the chosen neurons. Using the DAPI signal as a guide, the nuclei of these cells were traced and then a band expanding a uniform distance of 2 μm around each nucleus was drawn using the “draw band” function in ImageJ. Empirically, this band provides a close approximation of the cell soma boundary. Using ImageJ (NIH), the background signal was measured in three distinct places determined to lack true signal in each image, and the average mean background intensity was subtracted from the entire image. Finally, signal in the GAD65 channel was binarized, the adjustable watershed algorithm was applied, and GAD65 puncta within each band surrounding selected nuclei were quantified using the “analyze particles” function (size=0.1-10 and circularity=0.01-1.00). The number of puncta per band was divided by the area of each band to give a per soma density of GAD65 puncta. Between 16-40 cells were analyzed per slice. A subset of neurons in the CA1 region exhibit very high GAD65 immunofluorescence which fills the cell soma. Such cells were excluded from analysis.

### 2.8 Statistical analysis

Statistical analyses were performed using the R programming language (R Core Team, 2022) or SPSS. Linear mixed effects modeling was conducted in R utilized packages lmerTest (Kuznetsova, Brockhoff, & Christensen, 2017) and emmeans (Lenth, 2022). Bootstrapped multiple comparisons tests were conducted using custom written code in R. Specific statistical tests are described in the text and figure legends.

## 3. Results

### 3.1 Application of the extracellular domain of Sema4D to acute hippocampal slices decreases excitability

Previously we observed that the ECD of Sema4D fused to the Fc region of human IgG (Sema4D-Fc) mediates an increase in GABAergic synapse density as evidenced by immunostaining and seizure suppression in rodent hippocampus (Acker et al., 2018). Here, we sought to determine the effect and time-course of Sema4D treatment on the hippocampal circuit. We utilized a recording configuration and set-up that allows the recording of stimulation-evoked field hippocampal activity for long periods of time (> 12 hours; (Redondo et al., 2010). Recordings were performed in the CA1 region of acute hippocampal slices obtained from adult rats (aged postnatal day (P) 140-180). Evoked field responses comprise a positive-going field EPSP on which a negative-going population spike is superimposed. Following a stable 1-hour baseline, we bath-applied Sema4D-Fc or Fc control protein for 2 hours. A decrease in population spike amplitude was observed starting at ∼1 hour after Sema4D-Fc perfusion began and lasted throughout the recording period, including for 3 hours post-washout of Sema4D-Fc (Figure 1A). We quantified this effect at 3 hours post-washout and found a significant Sema4D-Fc-mediated decrease in population spike amplitude (Figure 1Bi & Ci).

**Figure 1.**
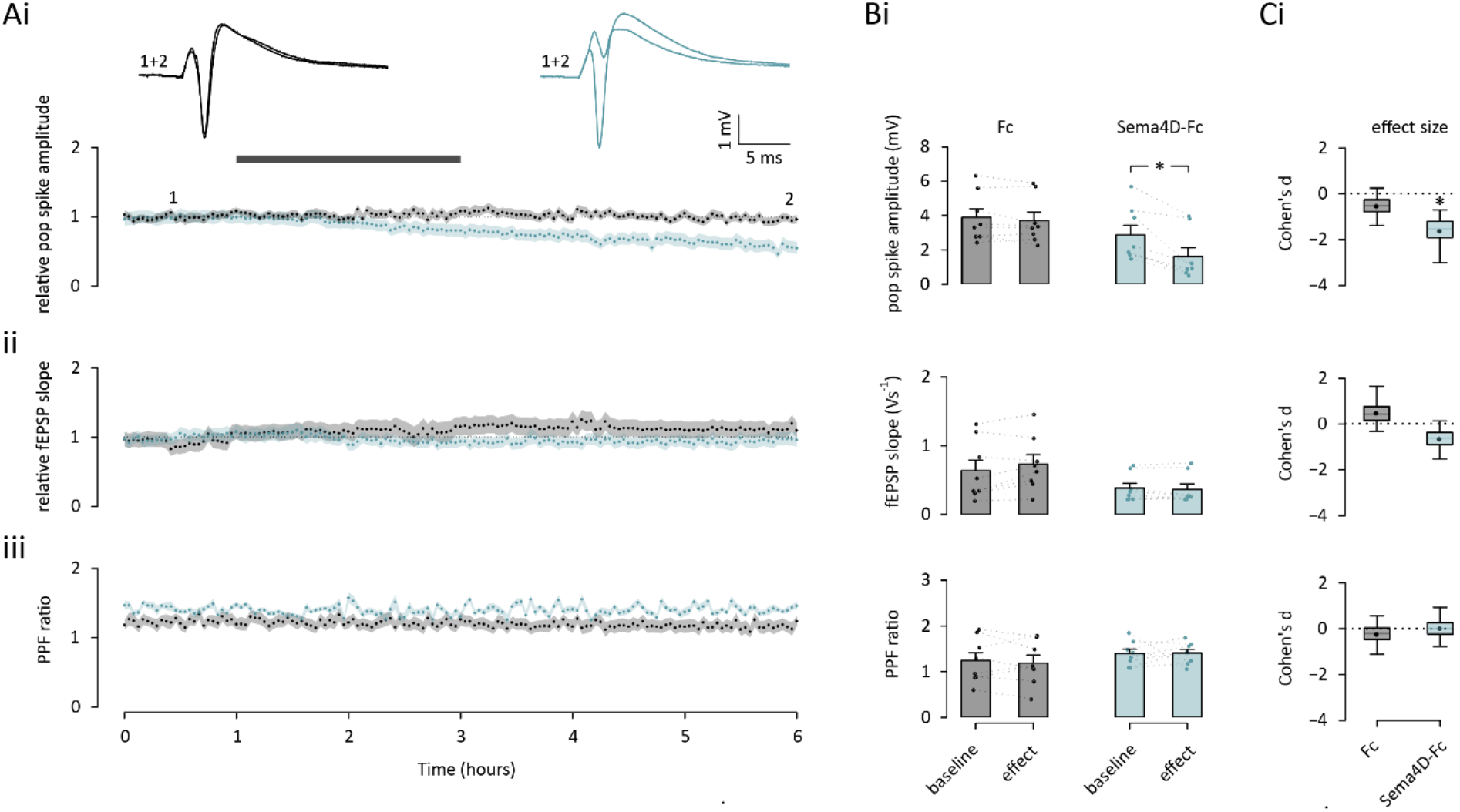
The extracellular domain of Sema4D protein decreases population spike amplitude in acute hippocampal slices. Sema4D-Fc treatment depressed population spike amplitude by ∼50% while having no effect on fEPSP slope or paired-pulse facilitation (PPF) ratio. Acute hippocampal slices were isolated from rats of both sexes at 20-25 weeks old. Responses were recorded by low-frequency stimulation of the Schaffer-collateral commissural pathway by stimulating electrodes placed in the stratum radiatum and recording responses at the CA1 pyramidal/stratum oriens border. A stable baseline (1 hour) was obtained before drug application for 2hrs. Comparisons were made 3hrs post washout (i.e., at t=6hrs). A, Time plots represent the pooled effects (both n=8) of control Fc. (black) vs Sema4D-Fc (blue) application to evoked population spikes (Ai), fEPSP slope (Aii) and PPF (measured as the ratio of fEPSP slopes to 2 successive pulses delivered at a 50ms interval; Aiii). Shaded areas are the (bootstrapped) s.e.m. Traces above the time plots represent the raw data comprising averages of time points within each experiment taken at the time points indicated. Fc / Sema4D-Fc application are shown in light gray. For both conditions, vertical and horizontal scale lines indicate 1mV and 5ms, respectively. B, Raw values (points) and summary statistics (mean (bars) and SEM (lines)) of population spike amplitude (Bi), fEPSP slope (Bii) and PPF ratio (Biii) shown at baseline and test time points as in A for control Fc (left, black) vs Sema4D-Fc (right, blue). In the group receiving Sema4D-Fc treatment, the mean spike amplitude at test was significantly lower than its baseline (bootstrapped multiple comparisons test with Holm-Šidák correction); no other conditions differed significantly from one another; * indicates alpha < 0.05. C, Box plot to illustrate Cohen’s d (bootstrap) for population spike amplitude (Ci), fEPSP slope (Cii) and PPF ratio (Ciii). Mean and median effect sizes are represented by point and line, respectively. The interquartile range and 95% confidence interval are illustrated by box and whiskers, respectively. Significant effects from raw data (B) are also illustrated on this plot by asterisk for convenience.

In addition, we failed to observe a change in initial slope of the fEPSP (Fig. 1Aii-Cii), a measure of L-glutamate mediated excitatory synaptic input, or paired-pulse facilitation (Fig. 1Aiii-Ciii), a measure of presynaptic release dynamics, in the Sema4D-Fc treatment condition. These data suggest that the depression of evoked firing observed upon Sema4D-Fc treatment is not due to a presynaptic effect which would dampen down excitatory synaptic transmission. However, we cannot definitely rule out an effect of Sema4D on neuronal intrinsic excitability. Nonetheless, given our previous studies demonstrating a rapid increase in GABAergic synapse formation in response to Sema4D application (Acker et al., 2018, Kuzirian, 2013 #2; Kuzirian et al., 2013), the most parsimonious explanation for the observed decrease in population spike amplitude is a Sema4D-dependent increase in evoked feedforward inhibitory transmission in the hippocampal slice.

### 3.2 Intrahippocampal infusion of the extracellular domain of Sema4D suppresses epileptiform activity and increases diazepam sensitivity in a SE model

Having confirmed a Sema4D-mediated decrease in evoked spiking activity in the adult hippocampus, we sought to determine if infusion of Sema4D-Fc into the hippocampus suppressed seizure activity and increased BZD sensitivity in the KA model of SE. We inserted a cannula bilaterally into the CA1 region of hippocampus in 10-12 week-old C57BL/6 male mice with hippocampal depth electrodes for EEG recordings affixed to their skulls. Two weeks later we infused 500nl of 100nM Sema4D-Fc or vehicle (Fc) via cannula over five minutes, waited one hour, and then induced seizures by a 20mg/kg intraperitoneal (i.p.) injection of KA (Figure 2A). Seizure activity was monitored throughout the experiment using EEG. Epileptiform activity was considered to be paroxysmal activity having a sudden onset and an amplitude at least 2.5x the standard deviation of the baseline and a consistent change in the Power of the fast Fourier transform of the EEG (Fig. 2B-C). In addition, SE was defined as persistent, unremitting epileptiform activity lasting longer than 5 consecutive minutes (Hooper et al., 2018; Sivakumaran & Maguire, 2016). All animals achieved SE by 1 hour post KA injection.

**Figure 2.**
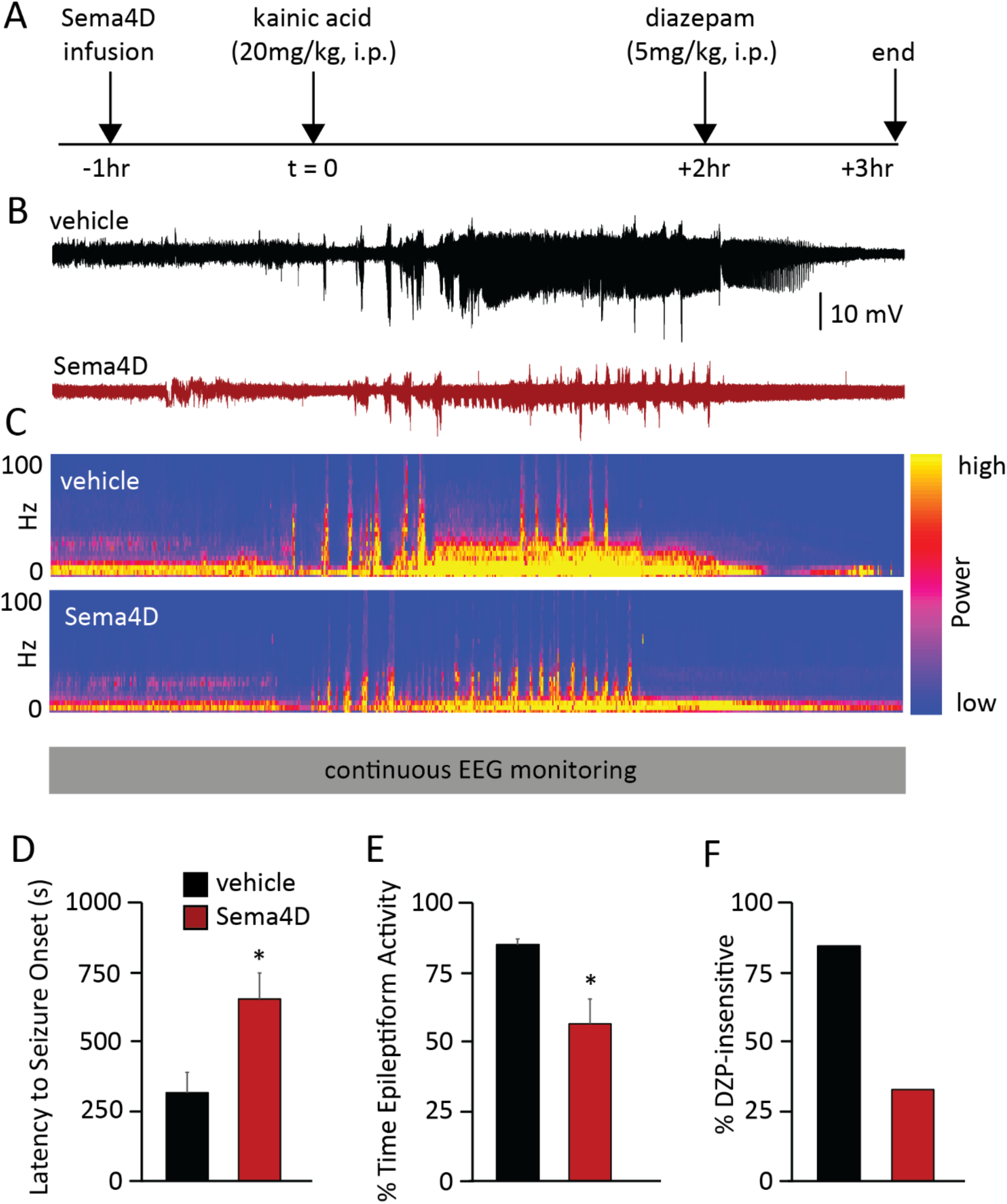
Sema4D treatment suppresses epileptiform activity and restores diazepam sensitivity *in vivo*. A, Experimental timeline B, representative electrographic seizure activity from vehicle (black) and Sema4D-Fc-treated mice (red) and C, representative EEG power spectra from vehicle (top) and Sema4D-Fc treated mice (bottom). C57BL/6 male mice aged 10-12 weeks were used for this experiment. Sema4D-Fc was infused bilaterally into the CA1 region of the hippocampus 1hr prior to administration of kainic acid (20mg/kg, i.p.). Electrographic seizure activity was recorded for the entire 4hr period of the experiment. For the two hours following KA injection, the following characteristics were quantified: D, latency to onset of the first seizure, E, cumulative epileptiform activity (% time) and F, percent of diazepam insensitive mice which was quantified from the one hour following diazepam injection (5mg/kg i.p.). n = 8 mice per experimental group; * denotes p<0.05 using a Student’s t-test.

Analysis of the two-hour recording period before diazepam treatment indicated that Sema4D-Fc infusion increased the latency to onset of the first seizure (Figure 2D). In addition, Sema4D infusion decreased the percentage of time the animals spent exhibiting epileptiform activity (Figure 2E), quantified as the cumulative time of all epileptiform activity during a 60-min recording period divided by 60 min, compared to control animals.

Two hours after KA injection, we delivered diazepam (5mg/kg) via i.p. injection and monitored the mice for one hour. We defined diazepam sensitivity as a reduction in the power of the electrographic signal and cessation of epileptiform activity as previously described (Hooper et al., 2018; Sivakumaran & Maguire, 2016). Mice were considered to be pharmacoresistant if diazepam treatment did not suppress SE within 10 mins of administration. Thus, by analyzing the EEG recordings obtained for one hour post-diazepam injection we observed that more animals in the Sema4D treatment group were sensitive to diazepam treatment compared to controls (Figure 2F). It is unlikely that this effect is due to decreased seizure burden in the Sema4D-Fc treated mice compared to control as all animals had reached SE before DZP was administered. Taken together, these data indicate that hippocampal infusion of soluble Sema4D-Fc suppresses KA-induced seizure activity and restores BZD sensitivity.

### 3.3 Adeno Associated Virus expressing Sema4D extracellular domain drives formation of inhibitory boutons in vitro

Since Sema4D treatment restores BZD sensitivity, we developed a method that would allow for more efficient delivery of the Sema4D protein. We began by asking if chronic, viral delivery of Sema4D could promote GABAergic synapse formation similarly to treatment with the purified protein. We created Adeno-Associated Viruses (AAV) serotype 9 expressing either the full-length Sema4D protein (AAV-Sema4D-FL) or its extracellular domain (AAV-Sema4D-ECD) and a control virus encoding full-length CD4 (AAV-CD4-FL), a transmembrane protein that has no effect on GABAergic synapse development (Raissi, Staudenmaier, David, Hu, & Paradis, 2013), or its extracellular domain (AAV-CD4-ECD) (Figure 3A). To selectively drive expression in neurons, all constructs are expressed under control of the human Synapsin I promoter.

**Figure 3.**
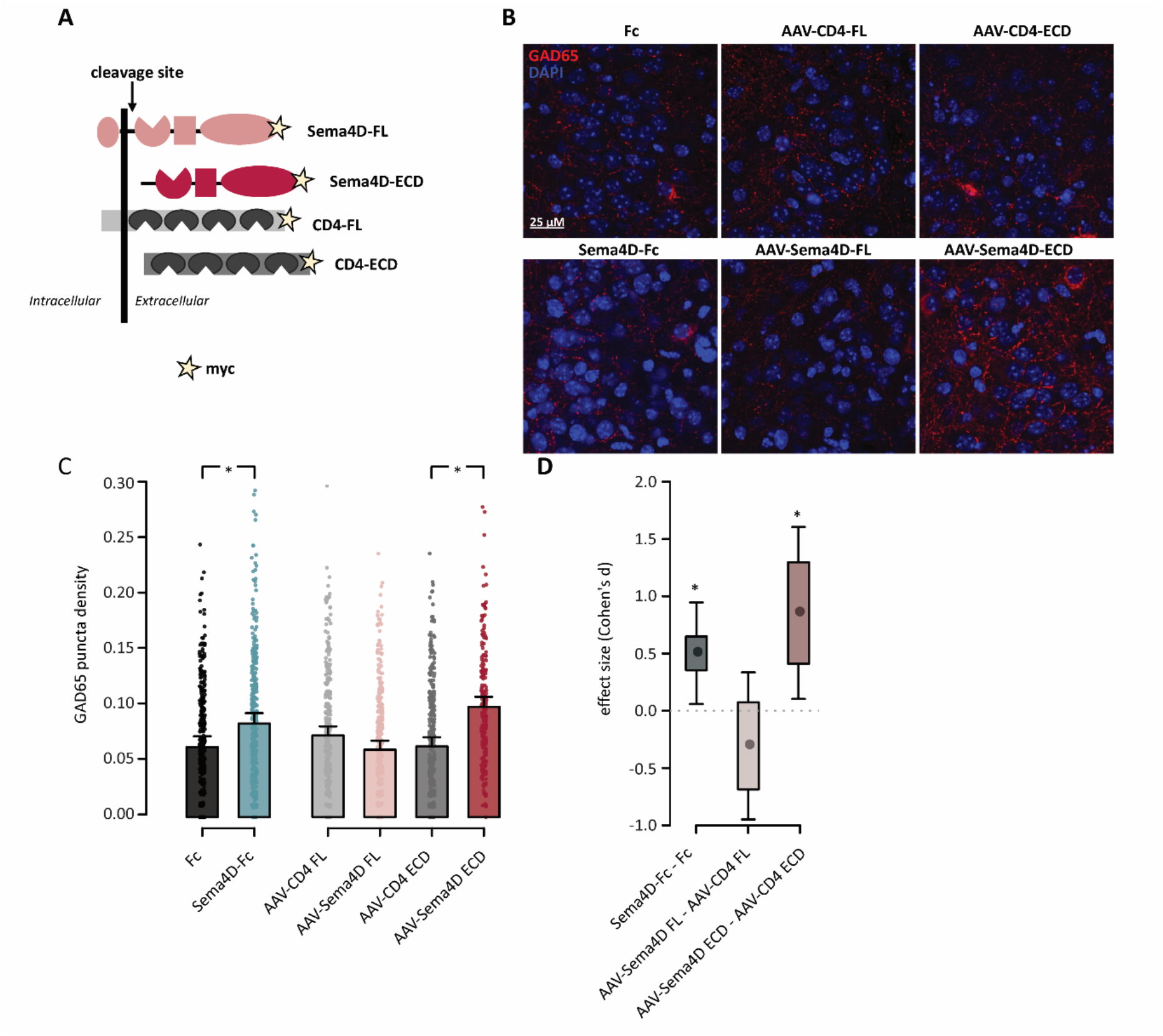
Validation of Sema4D-expressing adeno associated viruses *in vitro*. A, Schematic representing full-length or extracellular domain of Sema4D or CD4 protein encoded by the AAV viruses. B, Representative images of the CA1 principal cell layer in organotypic slices that were treated with Sema4D-Fc or Fc control proteins or infected with indicated AAV constructs; Sections are stained with an antibody that specifically recognizes GAD65 (red) and DAPI (blue). Scale bar represents 25μm. C, Density of GAD65 puncta per CA1 pyramidal cell soma; error bars are SEM. n = 233-351 neurons per treatment condition (represented by individual points), 16-40 neurons per slice, slices from 6-9 mice per treatment condition, 29 mice total. D, Box plot to illustrate the effect size, Cohen’s d. Mean effect size is represented by single points, interquartile range (IQR) and 95% confidence interval (CI) are illustrated by box and whiskers, respectively for the following comparisons: Fc vs Sema4D-Fc; AAV-CD4-FL vs AAV-Sema4D-FL; AAV-CD4-ECD vs AAV-Sema4D-ECD. A mixed effect linear model was fitted to the data with treatment as the fixed effect and animal and slice as random effects in order to control for variability that arises within animals and problems associated with pseudoreplication within slices (modeled as random intercepts) (Lazic, Mellor, Ashby, & Munafo, 2020). Consequently, data is presented as the estimated marginal means obtained from the model. Effect sizes (Cohen’s d), estimates of s.e.m, IQR, CI and multiple comparisons tests are based on these estimates. * denotes p<0.05 by conducting an ANOVA on the linear mixed effects model with subsequent multiple comparisons tests using a Tukey correction. Significant effects from raw data in C are also indicated by an asterisk in D for convenience.

As an initial test of efficacy of these viruses, we examined their ability to increase GABAergic synapse density in coronal, organotypic brain slices containing anterior hippocampus obtained from C57BL/6 mouse pups ages P6-P8. This time point was chosen for our initial verification because it is consistent with ours and others’ studies showing a pro-synaptogenic effect of purified Sema4D protein in *ex vivo* preparations (Acker et al., 2018; Frias et al., 2019; Kuzirian et al., 2013; McDermott, Goldblatt, & Paradis, 2018; Raissi et al., 2013). After 1 day in vitro (DIV1), slices were infected with a combination of AAV9.hsyn.GFP virus (Addgene) to express GFP and one of the Sema4D- or CD4-expressing viruses. At DIV6, slices were fixed and immunostained with an antibody that detects GAD65, an enzyme localized to the GABAergic presynaptic bouton. (Figure 3B). As a positive control for the function of Sema4D, we also infected a subset of slices with only AAV9.hsyn.GFP at DIV1 and treated with either Sema4D-Fc or Fc recombinant protein for two hours at DIV6 before processing the slices for immunostaining (Figure 3B).

We determined GABAergic synapse density by imaging the pyramidal layer in the CA1 region of hippocampus and quantifying the density of GAD65-positive boutons formed onto the cell soma of GFP+ neurons similar to our previous study (Acker et al., 2018). We found that neurons infected with AAV-Sema4D-ECD exhibited a significantly higher density of GAD65 puncta per soma as compared to the AAV-CD4-ECD control (Figure 3C, D). Notably, infection with AAV-Sema4D-FL did not increase the density of GAD65 puncta, perhaps indicating that soluble Sema4D-ECD is a more efficacious signaling moiety than membrane bound Sema4D. Importantly, the magnitude of change in GABAergic puncta density of ∼20-25% observed in this experiment is comparable to those observed using purified Sema4D-Fc protein in both this experiment (Figure 3C, D, Fc vs. Sema4D-Fc) and our previous experiments (Acker et al., 2018) see Figs. 1, 3, S2). Based on these results, we elected to test AAV-Sema4D-ECD in the KA model of SE, to determine whether chronic viral overexpression of Sema4D-ECD could reduce seizure symptoms and restore sensitivity to diazepam, similarly to infusion of purified Sema4D-Fc protein (Figure 2).

### 3.4 Delivery of AAV-Sema4D-ECD reduces epileptiform activity and restores BZD sensitivity in an SE model

We performed bilateral injections of either AAV-Sema4D-ECD or control AAV-CD4-ECD virus into the CA1 region of hippocampus into 10-12 week-old C57BL/6 male mice; we also affixed headmounts for EEG recording. Following a 1-week recovery period, we performed a 20mg/kg i.p. injection of KA to induce seizures and monitored seizure activity using EEG recording as described above (Fig. 4A-C). Analysis of the two-hour recording period after KA administration and before diazepam treatment indicated no difference between AAV-CD4-ECD and AAV-Sema4D-ECD virus in latency to onset of the first seizure (Figure 4D), which differs from our results with infusion of purified Sema4D-Fc protein (Fig. 2D above), and perhaps reflects differences in the levels of Sema4D-ECD at the time of seizure induction due to different delivery methods. However, we did observe that administration of AAV-Sema4D-ECD significantly reduced the amount of time spent in epileptiform activity as compared to mice treated with the AAV-CD4-ECD virus (Figure 4E), similar to the effect observed with Sema4D-Fc infusion (Fig. 2E), and consistent with an increase in inhibition in the hippocampi of these animals.

**Figure 4.**
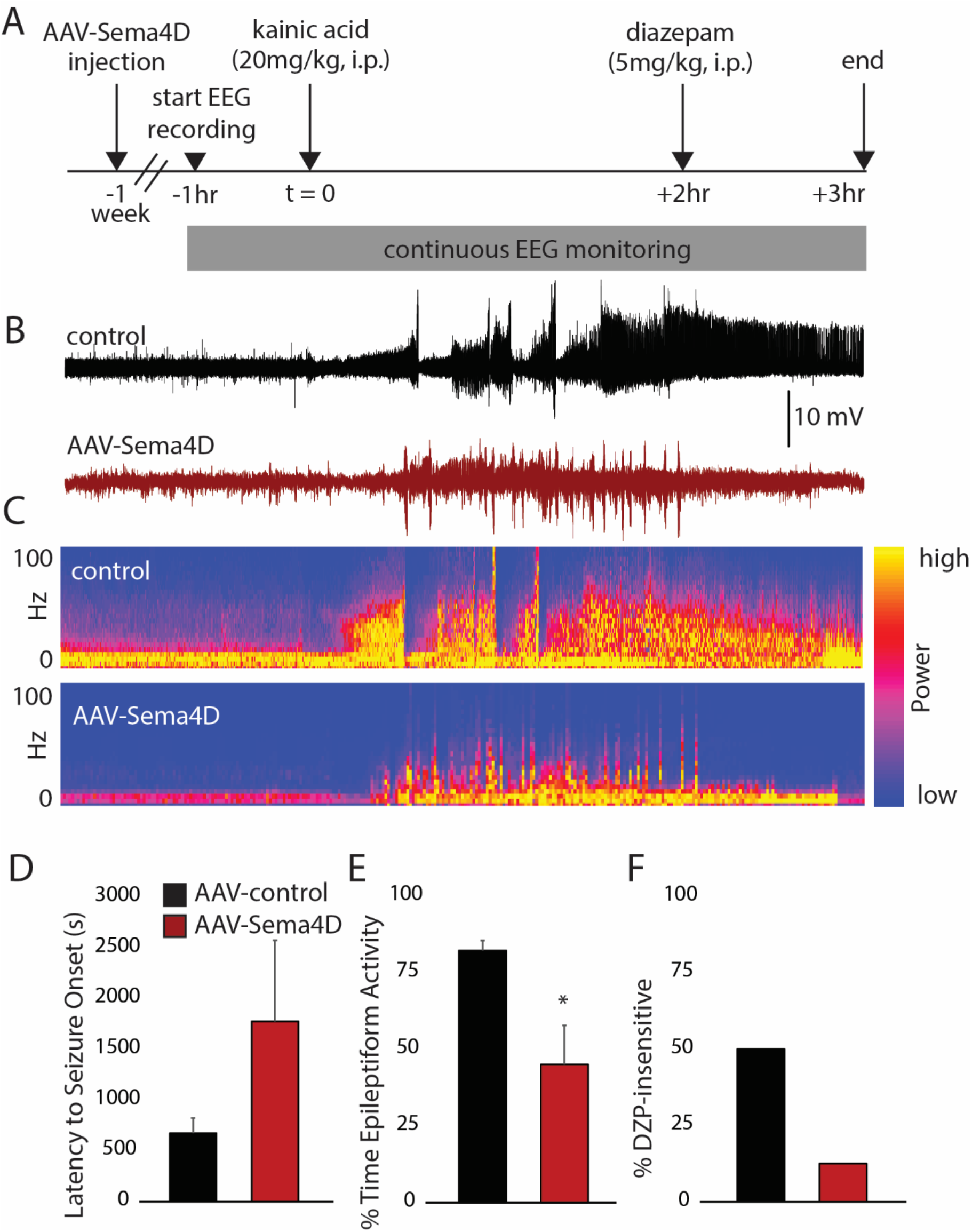
Delivery of AAV-Sema4D-ECD to hippocampus reduces epileptiform activity and restores BZD sensitivity. A, Experimental timeline B, representative electrographic seizure activity from control AAV-CD4-ECD (black) and AAV-Sema4D-ECD treated mice (red) and C, representative EEG power spectra from control AAV-CD4-ECD (top) and AAV-Sema4D-ECD treated mice (bottom). C57BL/6 male mice aged 10-12 weeks were used for this experiment. AAV-Sema4D-ECD or control virus was injected into the DG region of the hippocampus 1wk prior to administration of KA (20mg/kg, i.p.). Electrographic activity was recorded for 1 hr prior to and 3hrs after KA injection which includes the 1 hour post diazepam administration (5mg/kg, i.p.). For the two hours following KA injection, the following characteristics were quantified: D, latency to onset of the first seizure, E, cumulative epileptiform activity (% time) and F, percent of diazepam insensitive mice as quantified from the EEG recording for one hour following diazepam injection. n = 6-8 mice per experimental group; * denotes p<0.05 using a Student’s t-test.

Interestingly, in this experiment only 5 of 8 animals in the AAV-Sema4D-ECD group achieved SE by 1 hour post KA injection compared to 8 of 8 in the AAV-CD4-ECD cohort. Further, 2/8 control animals died compared to 0 for the AAV-Sema4D-ECD group. We speculate that these differences in severity of seizures post KA injection may be due to chronic expression of Sema4D in these animals, as opposed to the acute Sema4D-Fc treatment described above.

Two hours after KA injection, we delivered diazepam (5mg/kg) via i.p. and monitored the mice by EEG for an additional hour. Analysis of these recordings demonstrated that only 20% of AAV-Sema4D-ECD-infected mice that had achieved SE were BZD insensitive compared to 50% of control animals (Figure 4F), similar to our findings with infusion of purified Sema4D-Fc protein (Figure 2). In addition, DZP suppressed seizure activity in the 3 AAV-Sema4D-ECD-infected mice that failed to achieve SE. To validate the targeting of our viral injections to hippocampus, we performed post hoc immunostaining for GAD65 on animals that had been injected with AAV-Sema4D-ECD or AAV-CD4-ECD (Figure S1). We found an ∼ 20% increase in GAD65 immunostaining fluorescence intensity in the hippocampus of animals injected with AAV-Sema4D-ECD, consistent with correct targeting of the virus. Taken together, these results demonstrate that virally-mediated overexpression of Sema4D-ECD behaves similarly to infusion of purified Sema4D protein to increase inhibition and suppress seizures.

## 4. Discussion

In this study, we compared the efficacy of Sema4D treatment using two different methods: 1) an acute, intrahippocampal infusion of purified Sema4D protein via cannula and 2) a chronic, virus mediated delivery of Sema4D ECD. One week of chronic Sema4D overexpression ameliorated epileptic activity and increased BZD efficacy. The ECD of Sema4D is ∼700 amino acids and is not expected to cross the Blood-Brain Barrier (BBB). To overcome issues of BBB permeability, biotechnology and pharmaceutical companies are turning to strategies such as intrathecal injection of Adeno Associated Virus (AAV) to deliver therapeutics to the CNS (Mendell et al., 2021). Thus, our results support the translational potential of virally delivered Sema4D as an anti-seizure therapeutic for the treatment of both acute and chronic pharmacoresistant epilepsy.

The present study advances the case for Sema4D-dependent GABAergic synapse formation as a new avenue for developing anti-seizure therapeutics based on the following observations. First, we demonstrated that evoked CA1 excitability is suppressed by application of Sema4D protein to acute hippocampal slices and that this effect endures for at least 3 hrs, suggesting that Sema4D-mediated increased inhibition underlies the seizure suppression that we observe with *in vivo* application of Sema4D. Second, we showed that chronic delivery of Sema4D via viral transduction promotes increased GABAergic synapse density in *ex vivo* hippocampal organotypic slice cultures similar to purified Sema4D-Fc protein. Third, we demonstrated that acute application of Sema4D protein *in vivo* increases resistance to seizure and restores sensitivity to a BZD in the KA model of SE. Fourth, using the same KA model of SE, we discovered that chronic expression of the Sema4D-ECD in hippocampus *in vivo* is similarly efficacious to purified Sema4D protein in suppressing seizures and relieving BZD resistance.

The current study builds upon the following previous findings: 1) Sema4D-Fc protein treatment increases the density of functional, GABAergic synapses containing the BZD-sensitive gamma subunit of the GABA_A_ receptor in dissociated hippocampal neurons (Kuzirian et al., 2013), 2) Sema4D-Fc protein treatment increases the density of functional, GABAergic synapses in acute slice (Kuzirian et al., 2013), 3) Sema4D-Fc protein treatment stabilizes nascent, presynaptic GABAergic boutons in organotypic slice (Frias et al., 2019), 4) Sema4D-Fc infusion into the hippocampus of 6-8 week old mice both increases the density GAD65+ presynaptic termini in the hippocampus and suppresses seizures (Acker et al., 2018). All of these effects occur within a few hours of Sema4D treatment.

Taken together, these findings lead us to propose a model whereby Sema4D treatment promotes GABAergic synapse formation within 1 hour, resulting in a suppression of hippocampal network excitability, resilience to hyperactivity, and restored BZD sensitivity, presumably via incorporation of new GABAergic synapses into the hippocampus. Importantly, these data support the therapeutic potential of Sema4D in suppressing onset of SE and of its use as an “add-on” therapy for diazepam administration which is the current first line treatment for SE.

The amount of functional inhibition in a neuronal network is determined by every aspect of synapse biology: e.g., the number of receptors available in the membrane at the postsynaptic specialization, the number of active zones in the presynapse, and the number of synaptic contacts made between neurons. We interrogated the time course of the effect of Sema4D on population spike amplitude in CA1 in tissue taken from animals of an advanced age (Figure 1). The observation that Sema4D treatment begins to suppress evoked population spike amplitude within one hour of treatment, taken together with our immunohistochemistry data (Figure 3), (Acker et al., 2018; Kuzirian et al., 2013), suggest that Sema4D promotes the formation of functional GABAergic synapses resulting in decreased hippocampal excitability. These data also support our previous observations (Acker et al., 2018) that the cellular machinery which responds to Sema4D signaling to promote GABAergic synapse formation remains accessible in adult animals.

Studies in multiple model systems including from our lab have demonstrated that Semaphorins and their receptors are critical mediators of synaptogenesis (Ding, Oh, Sabatini, & Gu, 2011; Duan et al., 2014; Joo, Sweeney, Liang, & Luo, 2013; Mizumoto & Shen, 2013; O’Connor et al., 2009; Tran et al., 2009; Uesaka et al., 2014) Sema4D regulates GABAergic synapse formation via signaling through its high affinity receptor Plexin-B1 (Kuzirian et al., 2013; McDermott et al., 2018). It is widely accepted that Sema4D binding to Plexin-B1 causes dimerization and activation of the PlexinB1 intracellular GTPase activating protein (GAP) domain, triggering downstream signal transduction events that regulate the actin cytoskeleton and cell morphology (Li, Muller-Greven, Kim, Tamagnone, & Buck, 2021; Negishi, Oinuma, & Katoh, 2005; Oinuma, Katoh, & Negishi, 2004, 2006; Worzfeld et al., 2014).

Our current model of Sema4D signaling posits that Sema4D/Plexin-B1 signaling modulates the scaffolding machinery present at the synapse. Specifically, live-imaging of fluorescently-labeled synaptic proteins revealed that Sema4D signaling changes the subcellular localization of the GABAergic scaffolding protein gephyrin at the postsynapse (Kuzirian et al., 2013; Negishi et al., 2005). Rapid “splitting” of gephyrin-GFP puncta within ten minutes of Sema4D application resulted in additional gephyrin puncta which may nucleate new sites for GABAergic synapses to form. Furthermore, the c-Met receptor tyrosine kinase, which regulates actin depolymerization downstream of Sema4D/Plexin-B1 signaling (Swiercz, Worzfeld, & Offermanns, 2008), has been implicated in Sema4D-mediated stabilization of presynaptic GABAergic boutons and is specifically required in the presynaptic neuron for this effect (Frias et al., 2019). However, the precise signal transduction mechanisms underlying Sema4D/Plexin-B1 mediated GABAergic synaptogenesis are yet to be determined.

To design a more feasible route of Sema4D administration to the brain, we tested the ability of AAV9 virus expressing different portions of the Sema4D protein to promote GABAergic synapse formation in *ex vivo* organotypic slice cultures (Figure 3). Interestingly, while overexpression of Sema4D-ECD promoted GABAergic synapse development similarly to application of the purified Sema4D-Fc protein, overexpression of Sema4D-FL did not. Although the extracellular domain of Sema4D is cleaved from the neuronal cell surface, these results were unexpected as previous studies demonstrate that Sema4D can signal as a transmembrane bound or soluble extracellular domain (Raissi et al., 2013). It is plausible that differences in protein stability, expression, localization, or membrane insertion between Sema4D-ECD and Sema4D-FL may account for this disparity in synapse-promoting activity.

Both Sema4D and Plexin-B1 are expressed in inhibitory and excitatory neurons and glia in developing hippocampus (McDermott et al., 2018). Our functional studies show that Sema4D signals through Plexin-B1 expressed pre- and postsynaptically, suggesting that a bidirectional trans-synaptic signaling complex may regulate Sema4D-dependent GABAergic synapse formation. Since Sema4D-ECD expression was driven by a neuron-specific promoter in the AAV vector, our results demonstrate that overexpression of Sema4D-ECD in neurons is sufficient to increase the density of functional inhibitory synapses in hippocampus (Figure 4). We favor a model whereby virally-delivered Sema4D-ECD, secreted from neurons, acts in both an autocrine and paracrine manner to regulate this process.

Taken together our data suggest that Sema4D is uniquely suited to address problems of pharmacoresistance in SE, as Sema4D treatment in the hippocampus mitigates the BZD resistance observed in RSE, possibly by promoting GABAergic synapse development and/or stabilization. Irrespective of mechanism, our current data support further investigation of the translatability of Sema4D treatment for seizure suppression.

BZDs combat seizures and enhance inhibition by direct binding to GABA_A_Rs (see (Sills & Rogawski, 2020). However multiple groups showed that prolonged seizures are correlated with internalization or altered trafficking of GABA_A_Rs, with some studies citing decreased surface expression of the γ2 and β2/3 subunits of the GABA_A_R (Deeb et al., 2012; Goodkin, Joshi, Mtchedlishvili, Brar, & Kapur, 2008; Goodkin, Yeh, & Kapur, 2005; Naylor, 2005; Terunuma et al., 2008). Thus, therapeutics targeting SE that focus on methods to increase the density of GABA_A_Rs at the synapse in order to provide new sites for BZD binding could be effective as an add-on medication to BZD in RSE. Future studies will be required to determine whether Sema4D alters GABA_A_ receptor trafficking or localization.

The effect of ion flow through GABA_A_Rs can depolarize or hyperpolarize the neuron; this role changes throughout development and is mediated in large part by the constitutively active KCC2 and Cl^-^ homeostasis (Deeb et al., 2012). Thus, an alternative hypothesis regarding BZD resistance asserts that changes in KCC2 function interfere with BZD efficacy by altering the effect of net Cl^-^ flow across the cellular membrane, reversing the electrochemical driving force and the direction of GABA_A_-mediated current (Deeb et al., 2012; Moore, Kelley, Brandon, Deeb, & Moss, 2017). Interestingly, a recent study reported a proteomics approach which identified KCC2-interacting proteins. This study showed that gephyrin interacts with KCC2 and regulates its function by promoting KCC2 clustering at the membrane (Al Awabdh et al., 2022). It is possible that another mechanism by which Sema4D restores BZD sensitivity is via gephyrin splitting (Kuzirian et al., 2013) which could cause a redistribution of gephyrin and therefore KCC2 at the membrane. Future studies will be required to determine whether Sema4D alters GABAA receptor and/or KCC2 trafficking or localization. Intriguingly the same proteomics study that identified the KCC2-gephyrin interaction also identified the Sema4D receptor Plexin B1 as a putative interacting protein with KCC2 (Al Awabdh et al., 2022).

## 5. Conclusion

The number of patients suffering from pharmacoresistant epilepsy is expected to increase with the globally aging population. Alternative therapies to treat this disorder include resective surgery, deep-brain stimulation, and targeted laser ablation (Rai & Drislane, 2018). All have disadvantages: surgeries/ablation are destructive and invasive; open-loop stimulations are only palliative seizure-reducing therapies. Sema4D could be introduced by microinjection of protein for acute management of SE or as a gene therapy for chronic intractable epilepsy. The available evidence in rodent models of epilepsy suggests that, as a therapeutic, Sema4D has the potential to succeed where current AEDs fail. The ability of Sema4D to bypass mechanisms of pharmacoresistance combined with its potential to treat different seizure types, irrespective of etiology, in a minimally invasive fashion has the potential to be a disease modifying and life-altering therapy for the treatment of epileptic disorders.

## ETHICAL PUBLICATION STATEMENT

We confirm that we have read the Journal’s position on issues involved in ethical publication and affirm that this report is consistent with those guidelines.

## ACKNOWLEDGEMENTS

We thank all members of the Paradis lab and the Maguire lab for feedback and insight throughout the project.

