## Supplemental Figure 1 for "Semaphorin 4D induced inhibitory synaptogenesis restores benzodiazepine sensitivity in a mouse model of Status Epilepticus"

**Figure S1. Viral-mediated overexpression of Sema4D-ECD increases levels of a GABAergic synaptic marker in adult hippocampus**

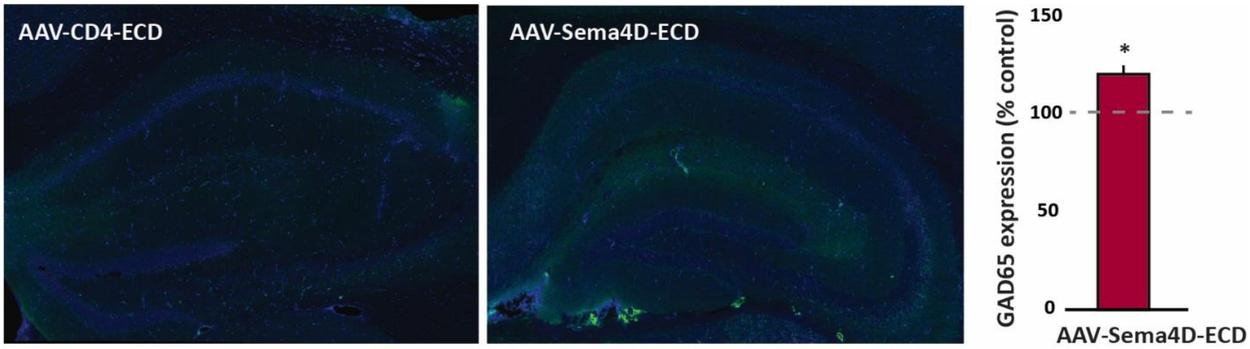


Animals expressing AAV-Sema4D-ECD or control virus used in experiments in Figure 4 were sacrificed at 2-3 weeks post virus injection and processed for immunostaining using an antibody that recognized GAD65. Imaging was performed on brain slices aligning with the site of injection. Representative images of GAD65 (green) and DAPI (blue) staining in the hippocampus 2-3 weeks following stereotaxic infusion of a control AAV-CD4-ECD virus or AAV-Sema4D-ECD. Regional intensity measurements in the hippocampus demonstrate an approximately 20% increase in GAD65 expression normalized to control slice (dotted line). n = 2-3 mice; 13-14 sections per experimental group; * denotes $p<0.05$ using a Welch's t-test.
